# SIRT7 links H3K36ac epigenetic regulation with genome maintenance in the aging mouse testis

**DOI:** 10.1101/2025.03.31.645534

**Authors:** Anna Guitart-Solanes, Mayra Romero, Irene Fernandez-Duran, Bryan A. Niedenberger, Cristina Madrid-Sandín, Ignasi Roig, Christopher B. Geyer, Alejandro Vaquero, Karen Schindler, Berta N. Vazquez

## Abstract

Reproductive aging is an increasing health concern affecting family planning and overall well-being. While extensively studied in females, the mechanisms driving male reproductive aging remain largely unexamined. Here we found that mammalian Sirtuin 7 (SIRT7) sustains spermatogenesis in an age-dependent manner through the control of histone 3 lysine 36 acetylation (H3K36ac). SIRT7 deficiency in mice resulted in increased levels of H3K36ac in spermatogonia and spermatocytes. In a germ cell line, SIRT7 deficiency disrupted nucleosome stability and increased vulnerability to genotoxic stress. Importantly, undifferentiated spermatogonia, which are required for continuous sperm production, decreased prematurely in *Sirt7^-/-^* mice and showed genome damage accumulation. These changes were concurrent with age-dependent defects in homologous chromosome synapsis and partial meiotic arrest. Taken together, our results indicate that SIRT7 connects H3K36ac epigenetic regulation to long-term genome stability in male germ cells, ensuring steady-state spermatogenesis during the lengthy male reproductive lifespan.

## INTRODUCTION

Reproductive potential declines with age in both female and male mammals. While the repercussions of advanced maternal age on mammalian oocyte quality have been extensively studied^1^, the impact of aging on male germ cell development have not received comparable attention. This lack of attention is due to the long- held view that aging exerted a minimal impact on male fertility. However, recent studies reveal that aging substantially influences male fecundity, impairing testis function and declining sperm quantity and quality^2–4^. Despite these important findings, the molecular mechanisms underlying age-related declines in male fertility remain poorly understood.

Spermatogenesis, the developmental program that continuously produces millions of sperm daily throughout the male reproductive lifespan, relies on functional spermatogonial stem cells (SSCs). SSCs tightly regulate a balance between self-renewal, to maintain the stem cell pool, and the production of transit-amplifying undifferentiated spermatogonia that subsequently commit to differentiation before entering meiosis^5^. During meiosis, spermatocytes complete a series of unique events, including double-strand break (DSB) formation, recruitment of DNA damage response (DDR) proteins, homology search, chromosome pairing, and, in some cases, resolution of DSB as crossover events^6^. Epigenetic changes are critical for multiple aspects of meiotic progression, including gene expression regulation and chromosome organization^7,8^. Defects in epigenetic events can result in meiotic arrest, leading to spermatogenic failure and male infertility^9,10^.

Sirtuins are an evolutionarily conserved family of NAD^+^-dependent protein deacylases and ADP- ribosyltransferases with major roles in aging and age-related diseases including cancer^11^, neurodegeneration^12,13^, and immunological disorders^14^. The mammalian genome encodes seven sirtuins (SIRT1-7) with differential cellular localization and functions. SIRT1, SIRT6 and SIRT7 primarily reside in the nucleus, where they promote genome homeostasis through epigenetic regulation^15^. SIRT2 predominantly resides in the cytoplasm but also binds to chromosomes in mitotic cells to ensure proper chromosome segregation^16^. SIRT3, SIRT4, and SIRT5 localize to mitochondria, where they regulate cellular metabolism and antioxidant defenses.

Sirtuins play major roles in mammalian reproduction by regulating a myriad of processes including germ cell development, meiotic recombination, chromosome segregation and redox homeostasis^17,18^. Increasing evidence links sirtuin function to reproductive aging, but the molecular mechanisms responsible for this association are poorly understood^18–20^. Female mice lacking *Sirt7* have reduced ovarian reserves and age- dependent subfertility characterized by compromised homologous chromosome synapsis that caused oocyte loss by birth and therefore a smaller primordial follicle pool^21^. Despite the importance of SIRT7 in normal early prophase I progression and reproductive longevity in females, its role in male gametogenesis has not been reported.

Here, we examined the reproductive consequences of *Sirt7* deletion in males and discovered an age-dependent fertility decline associated with premature testis degeneration, altered germ cell development, and compromised sperm DNA quality. Testes from young *Sirt7*^-/-^ males exhibited gene expression changes characteristic of premature aging and infertility that coincided with significant alterations in histone epigenetic signatures. In the absence of SIRT7, male germ cells had a constitutive increase in H3K36ac, with premeiotic spermatogonia and meiotic spermatocytes being the most affected populations. Lack of SIRT7 caused early onset of genome damage accumulation and loss of undifferentiated spermatogonia, and meiotic progression defects, suggesting an important role for SIRT7 in spermatogonia biology and early germ cell development. Taken together, our findings have major implications for male reproductive longevity and reveal a pivotal role for SIRT7 in safeguarding genome integrity by ensuring the proper establishment of epigenetic marks.

## RESULTS

### Loss of SIRT7 causes premature male reproductive defects

We previously reported an important role for SIRT7 in meiotic recombination in female mice, which is essential for the maintenance of a normal reproductive lifespan^21^. To assess SIRT7’s role in spermatogenesis and during aging, we analyzed reproductive parameters in previously validated *Sirt7* knockout (*Sirt7^-/-^*) male mice^22^. First, fertility studies were conducted by crossing wild-type (*Wt)* females with either *Wt* or *Sirt7^-/-^* males and numbers of litters and pups per litter were quantified (Fig. 1a-c). On average, despite variation in litter production among individuals, *Sirt7*^-/-^ males produced significantly fewer litters compared to *Wt* controls (Fig. 1b). Interestingly, although there was no significant difference in size of the first two litters, *Sirt7^-/-^* males produced fewer pups in subsequent litters, resulting in a reduction of the overall number of pups produced (Fig. 1c). In total, *Sirt7^-/-^*males sired nearly half as many pups as compared to similarly aged *Wt* control males. Together, our results reveal a significant age-dependent fertility decline in *Sirt7^-/-^* males.

**Figure 1.**
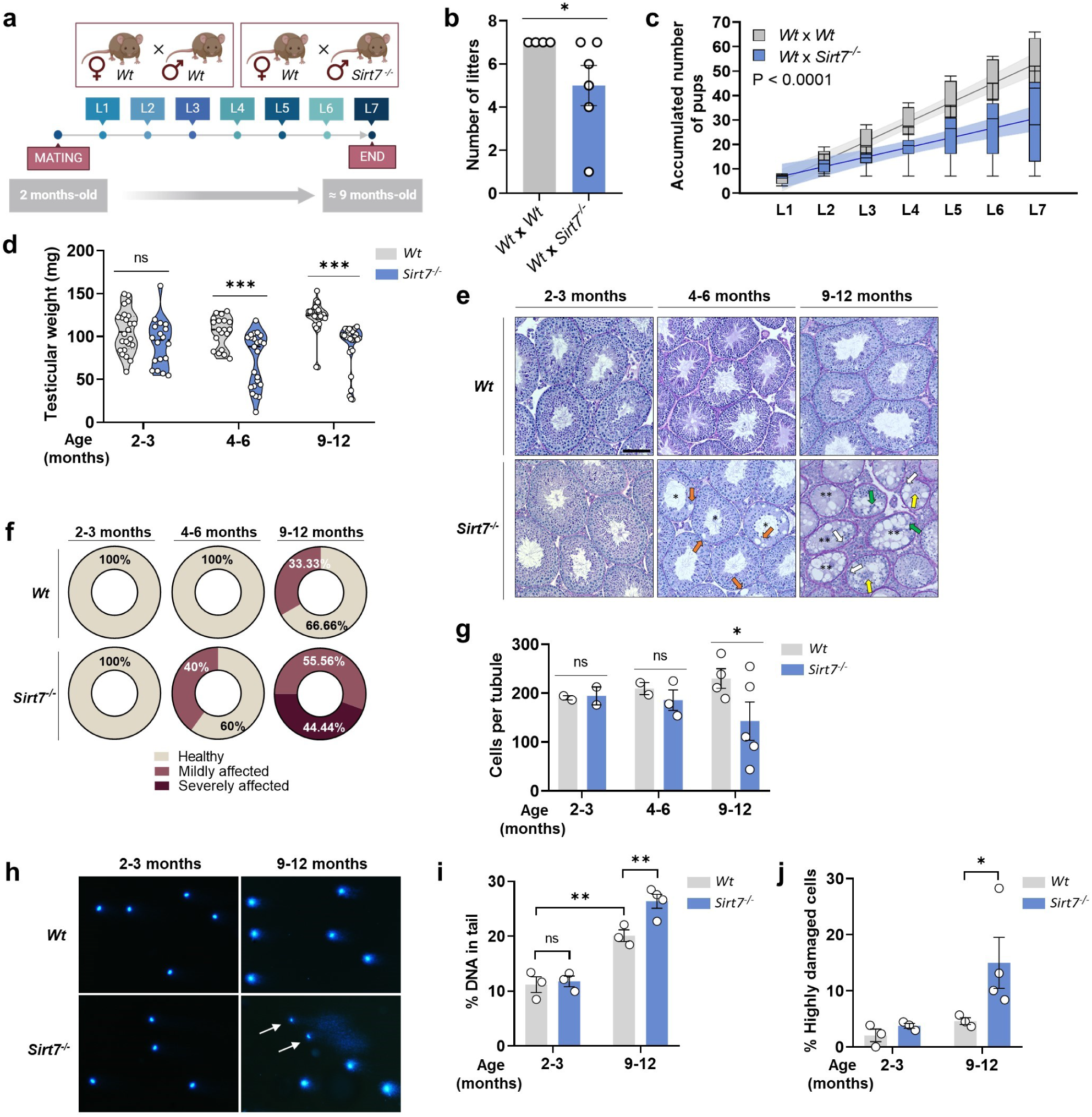
*Sirt7^-/-^* male mice display premature reproductive decline. **a)** Diagram of fertility trials. Single 2-month-old *Wt* female mice were caged with single 2-month-old *Wt* (n = 4) or *Sirt7^-/-^* (n = 6) mice. **b)** Numbers of litters produced by *Wt* x *Wt* and *Wt* x *Sirt7^-/-^* crosses. **c**) Accumulated number of pups produced during 7 litters (L1-L7) by *Wt* x *Wt* and *Wt* x *Sirt7^-/-^.* Each box shows the mean pup number of each litter. Linear regression analysis was performed to assess how different the tendencies between groups were, P < 0.0001. **d)** Testis weights of *Wt* and *Sirt7^-/-^* mice at different ages. n > 15 per group. **e)** Representative images of PAS-Hematoxylin-stained *Wt* and *Sirt7^-/-^* testes at different ages. Asterisks indicate seminiferous tubules with disrupted spermatogenesis; Arrows mark vacuolation initiation (orange), remaining spermatogonia (white), spermatocytes (green), and spermatids (yellow) in atrophic tubules. Scale bar = 100 µm. **f)** Pie charts indicating the percentage of animals presenting mild (>20% tubules with at least 1 vacuole) or severe (>20% tubules with abrogated spermatogenesis) histological affectation of the testes at the different age ranges. n = 3-9 testis samples per group. **g)** Quantitation of the number of germ cells per seminiferous tubule at each age range. Each data point represents one animal. **h)** Representative images of comet assays conducted in sperm. White arrows mark sperm with high DNA damage (‘hedgehog comets’). **i, j).** Graphs showing percentage of (i) DNA in comet tail and of (j) sperm with high DNA damage levels in 2-3- and 9-12-month-old *Wt* and *Sirt7^-/-^* sperm after comet assay. n = 3-4 sperm samples per genotype and age group, with > 200 cells quantified per sample.

Physiological testis aging is accompanied by histopathological changes, including impaired spermatogenesis and vacuolation due to germ cell losses within the seminiferous epithelium^3,23^. Notably, in 4- 6-month-old and older *Sirt7^-/-^* male mice, testis weights were significantly reduced, suggestive of testis degeneration (Fig. 1d). To visualize histopathological changes, Periodic Acid-Schiff (PAS)-Hematoxylin staining was performed on full testis sections from *Wt* and *Sirt7^-/-^* mice from three age groups (Fig. 1e). Mice at 2-3-months of age had normal-appearing spermatogenesis in both genotypes (Fig. 1e, f). In 40% of *Sirt7^-/-^* male mice at 4-6-months of age, abnormal vacuoles appeared in isolated seminiferous tubules, a phenotype also observed in one-third of testes from *Wt* mice at 9-12 months (Fig. 1e, f). This result revealed precocious degeneration and germ cell loss within seminiferous epithelia in *Sirt7^-/-^* mice. Interestingly, the observed age-related defects in *Wt* testes were consistent with a fertility decline with age in *Wt* males (Extended Data Fig. 1a). A much more severe vacuolation, characterized by complete depletion of germ cells in >20% of seminiferous tubules, affected ∼45% of 9-12-month-old *Sirt7^-/-^* testis. This effect was milder but still present (>20% tubules with ≥ 1 vacuole) in the remaining ∼55% of the aged testes (Fig. 1f). Consistently, the number of germ cells per seminiferous tubule were significantly reduced in testes from 9-12-month-old *Sirt7^-/-^* mice (Fig. 1g). Collectively, these data show that premature germ cell loss in *Sirt7^-/-^* testes underlies the age-related decline in the production of offspring by *Sirt7^-/-^*mice. Additionally, the variability in testis degeneration may explain the variation in numbers of litters produced by *Sirt7^-/-^* males.

Considering levels of DNA damage increase with age in both human and mouse sperm^23–25^, we compared DSBs levels using the comet assay in cauda epididymal sperm from *Wt* and *Sirt7^-/-^*mice of the 2-3-month-old and 9-12-month-old groups (Fig. 1h-j). Consistent with previous studies, our results revealed significantly increased sperm DNA fragmentation associated with aging, irrespective of genotype (Fig. 1i). Compared to *Wt* controls, *Sirt7^-/-^* sperm from 9-12-month-old mice had a significant increase in the percentage of DNA in comet tails (Fig. 1i). Importantly, ∼15% of sperm from 9-12-month-old *Sirt7^-/-^* mice exhibited high levels of DNA damage, demonstrated by their characteristic ‘hedgehog’ comet appearance (Fig. 1h, j). Collectively, these data indicate sperm from older *Sirt7^-/-^* mice have increased DNA damage, revealing a critical role for SIRT7 in maintaining genomic integrity to ensure the production of high-quality sperm during the aging process.

### SIRT7 absence results in transcriptomic changes resembling premature aging

SIRT7 acts as an important regulator of transcription in a gene context-dependent manner^26,27^. To define changes in the transcriptome of *Sirt7^-/-^* testes that could explain the age-related subfertility, we conducted bulk RNA-seq on whole testes. We aimed to identify the early transcriptomic changes underlying the onset of the premature reproductive aging in *Sirt7^-/-^* mice. For that, we compared testis from 2-3-month-old *Wt* and *Sirt7^-/-^*mice with those from 4-6-month-old *Wt* and *Sirt7^-/-^* mice, age at which the subfertility first appeared in *Sirt7^-/-^* mice (Fig. 2). Principal Component Analysis (PCA) revealed distinct PC dimensions in the *Sirt7^-/-^*testes compared to age-matched controls (Fig. 2a). Visualization of PCA-based unsupervised clustering showed an age-associated pattern in the first principal component (PC1), which accounts for 33% variance. These results suggest that 2-3-month-old *Sirt7^-/-^* mice transcriptionally resemble older *Wt* counterparts, pointing to early changes in gene expression in the absence of SIRT7 (Fig. 2a).

**Figure 2.**
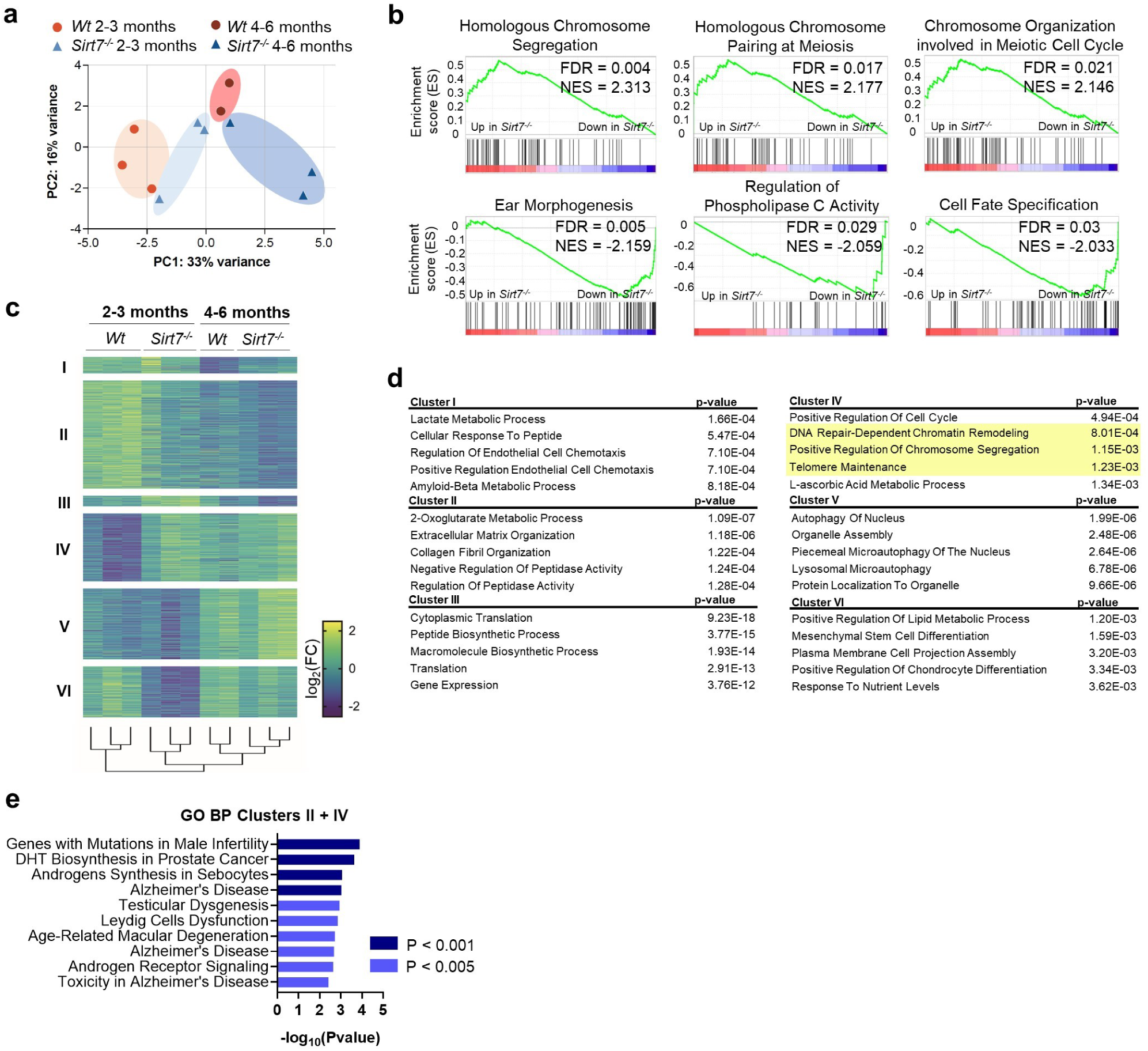
*Sirt7^-/-^* testes present a prematurely aged transcriptome with deregulation of transcripts related to chromatin organization. **a)** PCA of RNA-seq datasets derived from 2-3-month-old *Wt* (n = 3), 4-6-month-old *Wt* (n = 2), 2-3-month-old *Sirt7^-/-^* (n = 3) and 4-6-month-old *Sirt7^-/-^* (n = 3) testes. **b)** GSEA Enrichment plots of main gene signatures represented by upregulated (upper panels) and downregulated DEGs (lower panels) in 2-3-month-old *Sirt7^-/-^*samples compared to *Wt*. **c)** Heatmap of RNA-seq data showing all DEGs in at least one of the compared groups (P adj < 0.1) after an unsupervised clustering analysis. **d)** Top enriched signatures from the Elsevier Pathway Collection, corresponding to physiological pathways, for each of the gene clusters of the heatmap. **e)** Top enriched signatures from the GO Biological Process 2023 database for the genes in clusters II and IV of the heatmap (c).

Pairwise comparisons between genotypes identified differentially expressed genes (DEGs) between *Wt* and *Sirt7^-/-^* samples. In 2-3-month-old samples, 229 transcripts were significantly increased and 264 decreased in *Sirt7^-/-^*testes (Extended Data Fig. 2a). Gene Set Enrichment Analysis (GSEA) of upregulated DEGs in 2-3- month-old *Sirt7^-/-^* samples revealed gene signatures related to chromosome organization and meiosis (Fig. 2b), consistent with the known roles of SIRT7 in genome maintenance and meiotic progression^21,28^. Top downregulated gene signatures were related to morphogenesis, cell signaling and cell fate specification (Fig. 2b), uncovering novel potential roles for SIRT7 in male germ cell development.

Using unsupervised hierarchical clustering we identified 6 clusters regulated by either SIRT7, aging, or both (Fig. 2c). Gene ontology analysis (GO) of each gene cluster (Fig. 2d) revealed biological functions related to metabolism and extracellular matrix regulation (Cluster I, II and VI), protein synthesis (Cluster III), DNA damage repair and chromatin regulation (Cluster IV), and autophagy (Cluster V). Remarkably, Clusters II and IV contained prematurely downregulated and upregulated transcripts, respectively, in 2-3-month-old *Sirt7^-/-^* testes, showing high similarity to the levels in more aged *Wt* testes. Combined GO analysis of genes in Clusters II and IV revealed an association with age-related diseases and infertility signatures (Fig. 2e), suggesting their involvement in the premature reproductive decline of *Sirt7^-/-^*male mice. Importantly, several deregulated transcripts in 2-3-month-old *Sirt7^-/-^* testes included chromatin regulators, such as the histone methyltransferase suppressor of variegation 3-9 Homolog 2 (*Suv39h2*), the lysine-specific histone demethylase 4D (*Kdm4d*) and the Aurora A kinase (*Aurka*) (Extended Data Fig. 2b, c). These data reveal that, although testes of 2-3-month-old *Sirt7^-/-^*mice exhibited normal testicular architecture, the absence of SIRT7 triggered early transcriptional changes impacting chromatin regulation and long-term genome maintenance.

### SIRT7 regulates global H3K36ac levels in testis

Our transcriptomic analysis (Fig. 2) revealed that loss of SIRT7 resulted in altered transcript levels of genes encoding epigenetic regulatory proteins. Because SIRT7 is a bona fide epigenetic enzyme, we examined epigenetic changes in testes from 2-3- and 9-12-months-old *Wt* and *Sirt7^-/-^* mice (Fig. 3 and Extended Data Fig. 3). We first analyzed direct targets of SIRT7 deacetylase activity, including H3K18ac^26^ and H3K36ac^29^, by western blot analyses on whole testis lysates. Global levels of H3K18ac were similar in both *Wt* and *Sirt7^-/-^* testes (Fig. 3a-b), consistent with previous reports suggesting a role for SIRT7 in regulating this epigenetic mark at specific loci and excluding global H3K18ac regulation^22,26^. In contrast, a dramatic increase in H3K36ac was detected in *Sirt7^-/-^* testes, irrespective of age. In 2-3-month-old *Sirt7^-/-^* testes, there was a significant ∼6-fold increase in H3K36ac levels, as compared to a significant ∼4-fold increase in testes from older mice (Fig. 3a-b). Increased H3K36ac levels could preclude or limit deposition of H3K36 methylation^30^. However, despite major changes in H3K36ac in *Sirt7^-/-^*testes, global H3K36me2/me3 levels did not differ between genotypes (Fig. 3a-b). Further assessment of global levels of epigenetic marks linked either to spermatogenesis progression, such as H3K4 methylation^31^, or to sirtuin activity, such H4K20 and H3K9 methylations^16,32^, revealed no significant differences in *Sirt7^-/-^* testes (Extended Data Fig. 3a, b). Together, these results indicate SIRT7 is required to maintain deacetylated H3K36 in the mouse testis.

**Figure 3.**
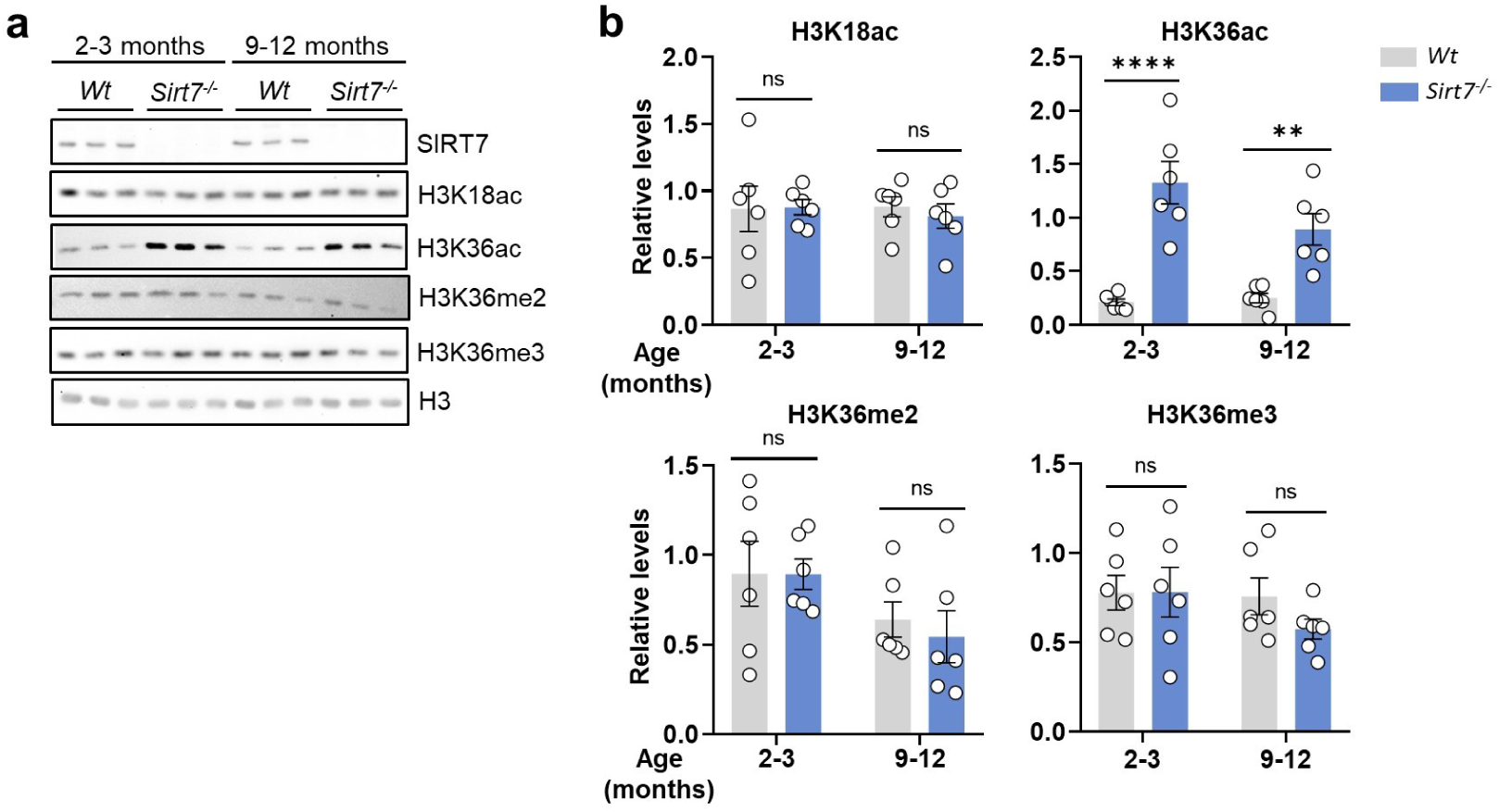
SIRT7 regulates H3K36ac in mouse testes. **a)** Western blots of modified histones in whole testis protein lysates from 2-3- and 9-12-month-old *Wt* and *Sirt7^-/-^* mice. **b)** Densitometry-based quantifications of the relative levels of the epigenetic marks. Levels of each epigenetic mark were normalized to H3 levels. n = 6 testis samples per genotype and age group.

### SIRT7 regulates H3K36ac in spermatogonia and spermatocytes

Western blot analyses showed a clear deregulation of H3K36ac levels in SIRT7-deficient testes (Fig. 3). Analyses of publicly available scRNA-seq data from human^33^ and mouse^34^ testes indicated that *Sirt7* transcript levels were abundant in spermatogonia and spermatocytes but substantially decreased in post-meiotic spermatids (Fig. 4a, b), suggesting a previously unrecognized role for SIRT7 in early spermatogenic populations. To understand the contribution of SIRT7 and H3K36ac in spermatogenesis during testicular aging, we analyzed H3K36ac distribution in testicular sections from 2-3- and 9-12-month-old *Wt* and *Sirt7^-/-^* mice (Fig. 4c). Independently of age, absence of SIRT7 led to a marked increase in H3K36ac across all germ cell populations, with premeiotic spermatogonia and meiotic spermatocytes displaying the largest fold changes (Fig. 4 c-e). Notably, in *Wt* testes we observed an inverse correlation between *Sirt7* and H3K36ac levels, with H3K36ac levels barely detectable in spermatogonia and spermatocytes and increasing in round spermatids (Fig. 4d, f), while *Sirt7* mRNA levels display the opposite trend (Fig. 4g). Levels of H3K36ac remained high in elongating spermatids, but as expected were depleted in condensing spermatids, in which the majority of histones are replaced by protamines (Extended Data Fig. 4a). The increase in H3K36ac and the decline in *Sirt7* levels from spermatogonia to spermatids points to an important role of SIRT7-dependent H3K36ac deacetylation in this developmental process. Notably, *Sirt7* transcript levels were the highest among all sirtuins in human spermatogonia and spermatocytes and were also among the highest in these populations in mice (Extended Data Fig. 4b-d), reinforcing the potential role of SIRT7 in early spermatogenesis.

**Figure 4.**
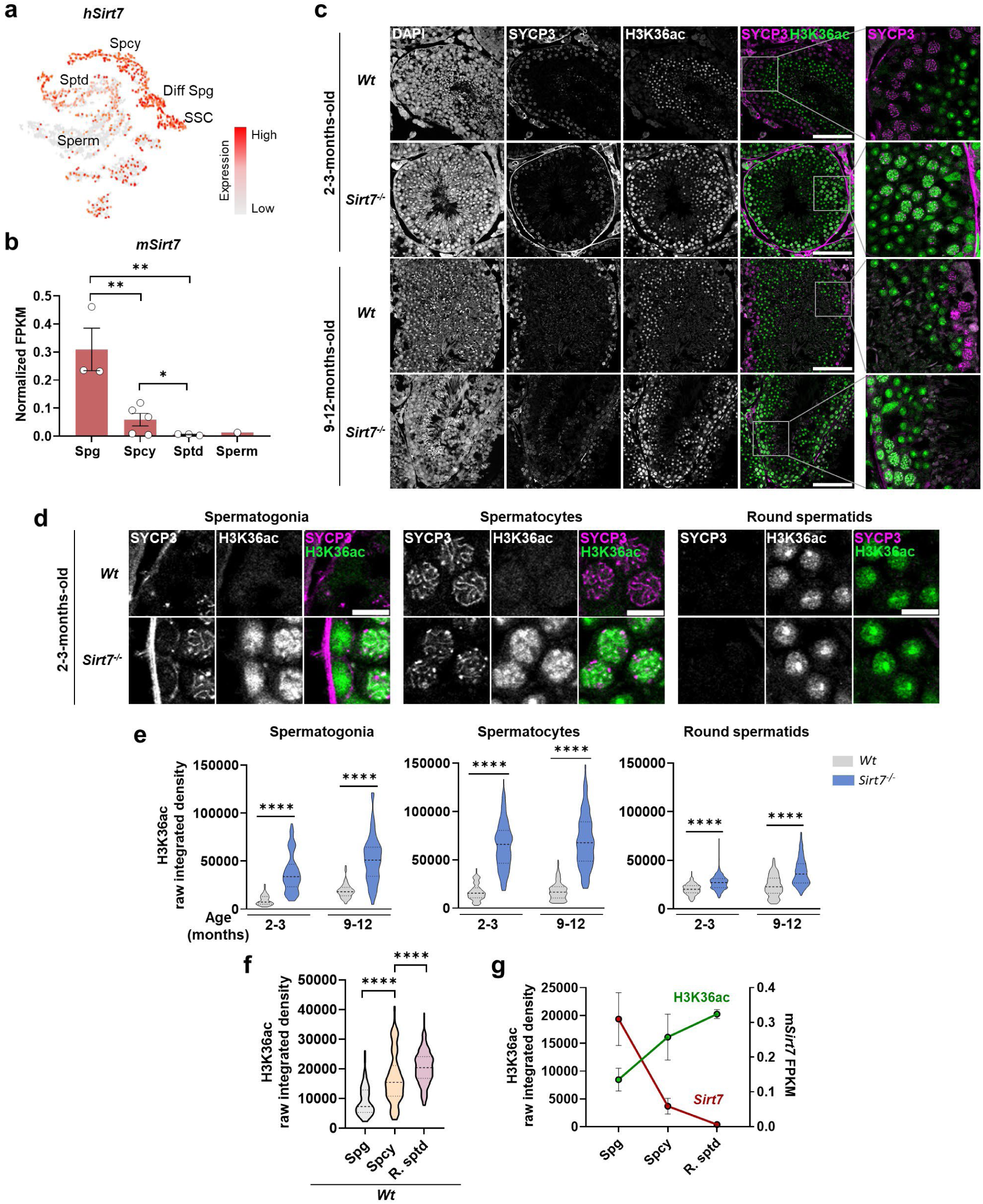
SIRT7 regulates H3K36ac in spermatogonia and spermatocytes. **a)** tSNE plot of single-cell transcriptome data from young human testes showing the expression levels of *Sirt7* in different testis cell types. **b)** Data from scRNA-seq of mouse testes showing the transcript levels of mouse *Sirt7* in male germ cell types. **c)** Immunostaining of SYCP3 and H3K36ac in testis sections of 2-3- and 9-12- month-old *Wt* and *Sirt7^-/-^* mice. Scale bar = 100 µm. **d)** Representative images of the immunostaining of H3K36ac in *Wt* and *Sirt7^-/-^*samples in spermatogonia, spermatocyte, and round spermatid cell populations. Spermatogonia were identified by their position near the basal membrane of the tubule, absence of SYCP3 staining, and characteristic morphology. Spermatocytes were identified by the presence of SYCP3 staining. Round spermatids were identified by absence of SYCP3 staining, adluminal position in the tubule, small size, and presence of a distinct chromocenter in DAPI staining. Scale bar = 10 µm. **e)** Quantification of H3K36ac fluorescence intensity in the different spermatogenic cell types in 2-3- and 9-12-month-old *Wt* and *Sirt7^-/-^* samples. **f)** Quantification of H3K36ac fluorescence intensity in the different spermatogenic cell types in 2-3- month-old *Wt* samples. **g)** Correlation graph of H3K36ac fluorescence intensity in 2-3-month-old *Wt* with mouse *Sirt7* transcript levels from public datasets in the indicated germ cell populations. All quantifications were made with n = 3 different animal samples for each age and cell group with >50 cells quantified for spermatogonia and >200 for the other groups. In this figure: SSC, Spermatogonial stem cells; Diff Spg, differentiating spermatogonia; Spg, spermatogonia; Spcy, spermatocytes; Sptd, spermatids; R. Sptd, round spermatids.

We further analyzed H3K36ac dynamics during prophase I in meiotic chromatin spreads from 2-3- and 9- 12-month-old *Wt* and *Sirt7^-/-^*mouse testes. *Wt* spermatocytes exhibited low H3K36ac levels in leptonema and zygonema that increased in pachynema and to a higher extent in diplonema (Extended Data Fig. 4e, f). Consistently, *Sirt7* transcript levels declined during meiosis (Extended Data Fig. 4g).

### Lack of SIRT7 results in altered chromatin accessibility and DNA damage accumulation

Previous work showed that H3K36ac facilitates chromatin relaxation in yeast^30^, raising the possibility that SIRT7-dependent H3K36ac activity in spermatogonia and spermatocytes regulates chromatin accessibility. To test this possibility in a readily manipulable cell population of germ cell origin, we introduced a CRISPR-Cas9- induced deletion of the *Sirt7* gene into the spermatocyte-derived immortalized mouse cell line GC-2spd(ts). This deletion led to high H3K36ac levels as in primary cells (Fig. 5a). Micrococcal Nuclease (MNase) assays were then conducted with *Wt* and *Sirt7^-/-^* cells, consisting of MNase treatment for increasing time periods, followed by DNA fragment analysis (Fig. 5b-d). Notably, the total amount of DNA remaining after digestion was significantly reduced in the absence of SIRT7 (Fig. 5b, c). Chromatin treatment with MNase resulted in linker DNA digestion and mononucleosome accumulation, as observed in *Wt* cells (Fig. 5b, d). Remarkably, mononucleosome accumulation did not occur in *Sirt7^-/-^*cells, but instead a depletion of this fraction was observed (Fig. 5d). Our results suggest SIRT7 deficiency disrupts chromatin structure, possibly affecting the nucleosome core organization.

**Figure 5.**
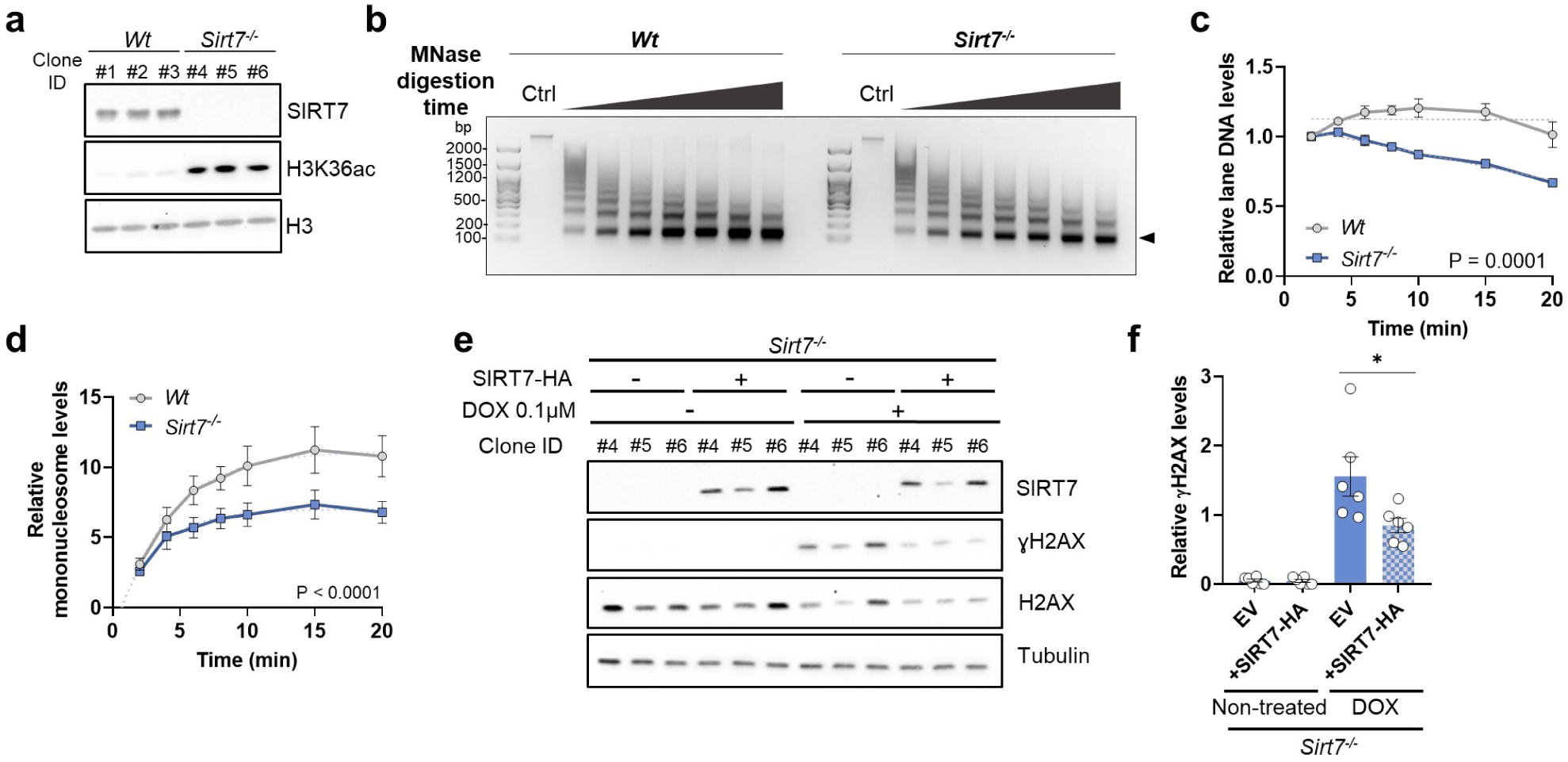
Impaired chromatin conformation and DNA damage accumulation in the absence of SIRT7 in a spermatocyte-derived cell line. **a)** Western blot of SIRT7 and H3K36ac levels in three *Wt* and three *Sirt7^-/-^* GC-2spd(ts) CRISPR-Cas9- generated clones. **b)** Agarose gel banding showing digestion of the chromatin of *Wt* and *Sirt7^-/-^*GC-2spd(ts) cells in the presence of MNase over increasing time periods (0, 2, 4, 6, 8, 10, 15 and 20 min). Black arrowhead at the left of the image marks the bands corresponding to the mononucleosome fraction. **c)** Densitometry-based quantification of chromatin digestion showing total amount of DNA at each time point normalized by the amount of DNA at t = 0 min. Linear regression analysis was performed to test how different the digestion tendencies were between genotypes, P = 0.0001. n = 3 independent clones per genotype. **d)** Densitometry-based quantification of chromatin digestion showing the intensity of the mononucleosome fraction normalized by the amount of DNA at t = 0 min. Non-linear regression analysis was performed to test how different the digestion tendencies were between genotypes, P < 0.0001 **e)** Western blot of SIRT7 and ɣH2AX in non-treated and 0.1 µM doxorubicin (DOX)-treated *Sirt7^-/-^* GC-2spd(ts) cells after the reintroduction of EV or SIRT7-HA in *Sirt7^-/-^* GC-2spd(ts) cells (3 independent clones per condition). **f)** Densitometry-based quantification of ɣH2AX levels relative to H2AX levels of experiment (e), n = 2 replicates.

Because chromatin structure is important for DNA damage repair and genome maintenance, we assessed the extent to which *Sirt7^-/-^* GC-2spd(ts) cells were susceptible to DNA damage accumulation as compared to *Wt* controls. Under basal conditions, *Wt* and *Sirt7^-/-^* GC-2spd(ts) cells had similarly low levels of phosphorylated H2AX (γH2AX,) a histone mark associated with DSB formation^35^ (Extended Data Fig. 5a, b). As expected, treatment with the genotoxin doxorubicin increased γH2AX signal accumulation in both genotypes but, importantly, *Sirt7^-/-^* cells showed higher levels of γH2AX compared to *Wt* controls (Extended Data Fig. 5a, b). Additionally, reintroduction of SIRT7 in *Sirt7^-/-^* GC-2spd(ts) cells significantly reduced γH2AX levels (Fig. 5e, f). Together, our results link SIRT7 activity to changes in chromatin accessibility and the prevention of DSB accumulation upon environmental stress.

### SIRT7 absence results in DNA damage accumulation in undifferentiated spermatogonia

*Sirt7* transcript levels are high in spermatogonia (Fig. 4a, b), the cells that provide the foundation for the continuous supply of spermatogenic cells throughout the reproductive lifespan. Given the critical role of SIRT7 in chromatin regulation and genome maintenance, we hypothesized that SIRT7 activity sustained genome integrity in spermatogonia. We first identified undifferentiated spermatogonia^5^, by immunostaining for PLZF in both 2-3-month-old and 9-12-month-old *Wt* and *Sirt7^-/-^*testes. Analysis of PLZF^+^ cells in 2-3- month-old *Wt* and *Sirt7^-/-^*testis sections revealed no significant changes in undifferentiated spermatogonia numbers (Fig. 6a, b). In 9-12-month-old testes, however, there was significant decline in undifferentiated spermatogonia numbers per tubule in *Sirt7^-/-^* testes compared to *Wt* controls (Fig. 6a, b). Further analysis of DSB accumulation in *Wt* and *Sirt7^-/-^* testis samples was conducted by co-immunostaining for γH2AX and PLZF. The results revealed significant accumulation of γH2AX in undifferentiated spermatogonia with age, which was more pronounced in *Sirt7^-/-^*cells (Fig. 6c, d). Collectively, these findings highlight an important role for SIRT7 in maintaining undifferentiated spermatogonia population during aging.

**Figure 6.**
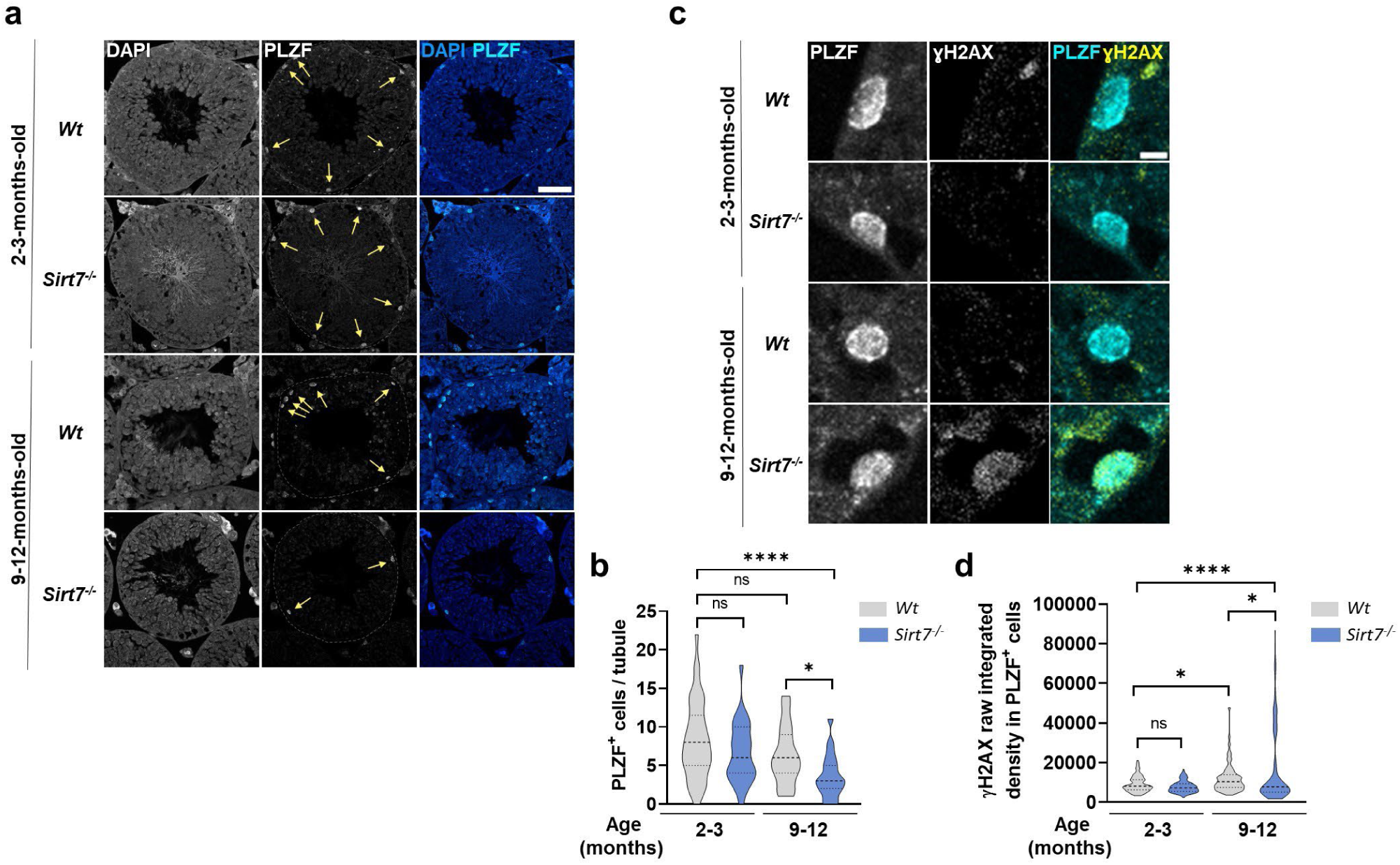
Lack of SIRT7 leads to loss of undifferentiated spermatogonia with aging linked to premature DNA damage accumulation. **a)** Immunostaining of PLZF in testis sections of 2-3- and 9-12-month-old *Wt* and *Sirt7^-/-^* mice. Dotted lines delimit the area of the seminiferous tubules. Yellow arrows mark PLZF^+^ undifferentiated spermatogonia. Scale bar = 50 µm. **b)** Quantification of the number of PLZF^+^ undifferentiated spermatogonia per seminiferous tubule. n = 3 animal samples per genotype and age group, >10 tubules analyzed per sample. **c)** Co-immunostaining of PLZF and ɣH2AX in testis sections of 2-3- and 9-12-month-old *Wt* and *Sirt7^-/-^* mice. Image of 9-12-month-old *Sirt7^-/-^* undifferentiated spermatogonia is representative of the cells with high ɣH2AX levels in that group. Scale bar = 5 µm. **d)** Quantification of ɣH2AX fluorescence intensity in PLZF^+^ undifferentiated spermatogonia.

### SIRT7 absence impairs meiotic progression in an age-dependent manner

Given the premature decline and increased DSB accumulation in 9-12-month-old *Sirt7^-/-^* undifferentiated spermatogonia, we investigated the extent to which genome instability persisted in early stages of meiotic progression. Using co-immunostaining for γH2AX along with SYCP3, a lateral element protein of the synaptonemal complex^36^, we identified and quantified the distinct prophase I stages in *Wt* and *Sirt7^-/-^* testes. In *Wt* spermatocytes, γH2AX signals were dispersed during leptonema and zygonema, coinciding with DSB formation and the onset of chromosome pairing. At the completion of chromosome synapsis in pachynema, γH2AX disappeared from autosomes and remained only in the XY bivalent, also called the sex body, where it promotes meiotic sex chromosome inactivation^37^ (Extended Data Fig. 6a). Chromatin spreads from 2-3-month- old *Wt* and *Sirt7^-/-^* testes had similar percentages of leptotene, zygotene, pachytene, and diplotene spermatocytes (Fig. 7a). In contrast, 9-12-month-old *Sirt7^-/-^*testes had increased numbers of spermatocytes with pachynema- like SCYP3 axes but with persistent γH2AX signals (Fig. 7b-d). Further analyses of pachytene-like spermatocytes in aged *Sirt7^-/-^*samples revealed increased levels of HORMAD1, a protein that binds to unsynapsed chromosome axis (Fig. 7e, f), and an increase in RAD51 recombinase foci (Fig. 7g, h), linking the loss of SIRT7 to an age-dependent increase in homologous chromosome synapsis defects, defective DNA damage repair and impaired meiotic progression (Fig. 7i). Overall, our results reveal a critical role for SIRT7 as a key regulator of chromatin structure and genome stability in both spermatogonia and spermatocytes, with important consequences for male reproductive longevity.

**Figure 7.**
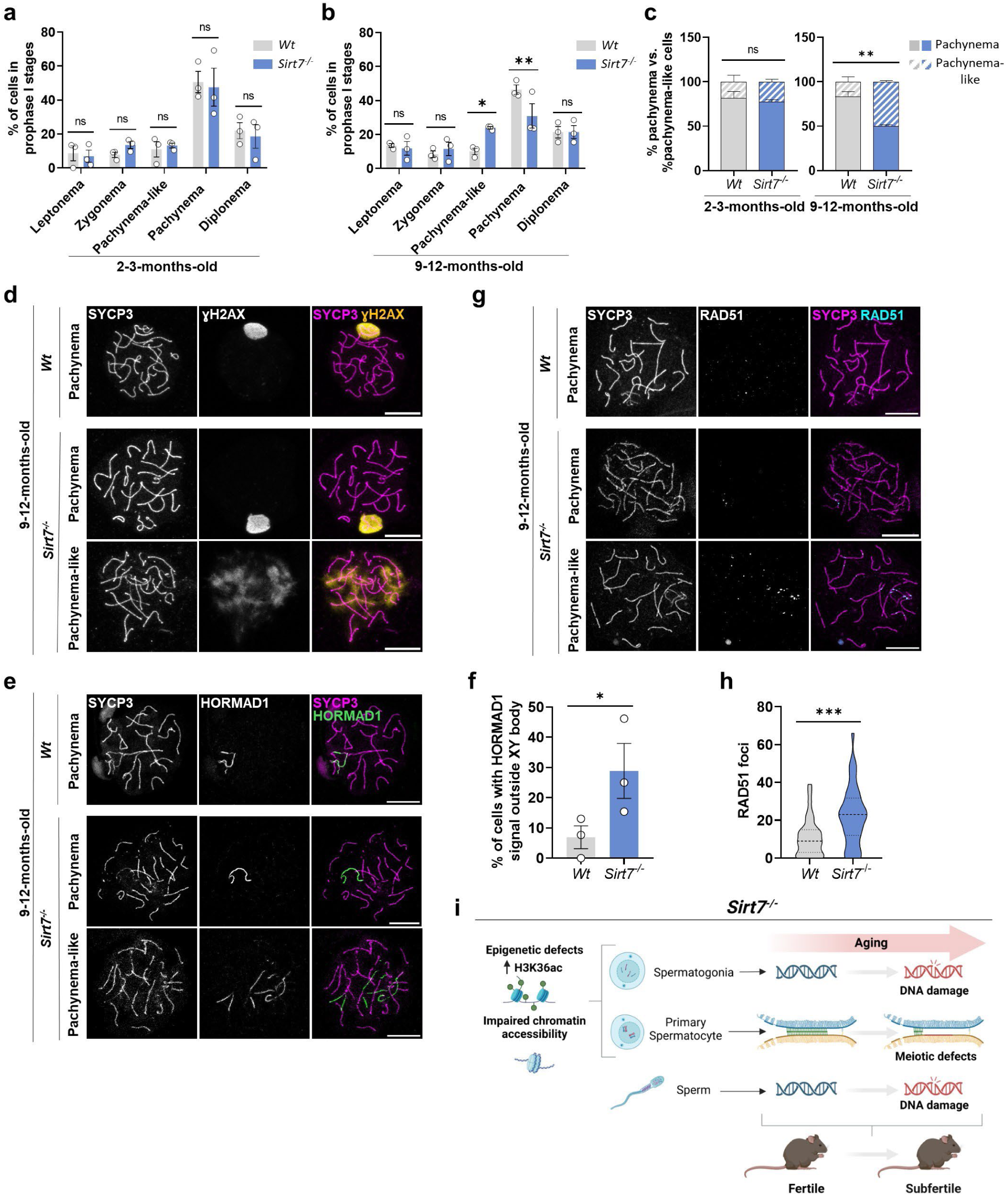
SIRT7 deficiency results in meiotic defects appearing with aging. **a, b)** Percentages of cells in the different meiotic prophase stages in chromosome spreads from (a) 2-3- and (b) 9-12-month-old *Wt* and *Sirt7^-/-^* testes. Each data point represents one animal. 200 cells were counted in each sample. **c)** Percentage of healthy pachynema cells and pachynema-like cells in chromosome spreads of 2-3- and 9-12-month-old *Wt* and *Sirt7^-/-^* testes. **d)** Representative images of SYCP3 and ɣH2AX-immunostained *Wt* healthy pachynema spermatocytes and *Sirt7^-/-^* pachynema and pachynema-like spermatocytes. Scale bar = 10 µm**. e)** Immunostaining of SYCP3 and HORMAD1 in *Wt* and *Sirt7^-/-^* pachynema and pachynema-like spermatocytes from 9-12-month-old *Wt* and *Sirt7^-/-^* mice. Scale bar = 10 µm**. f)** Quantification of the percentage of pachynema cells that present HORMAD1 signal outside the XY bivalent. Each data point represents one animal. **g)** Immunostaining of SYCP3 and RAD51 in 9-12-month-old *Wt* and *Sirt7^-/-^* pachynema and pachynema-like cells. Scale bar = 10 µm**. h)** Quantification of RAD51 foci number in *Wt* and *Sirt7^-/-^* pachynema and pachynema-like spermatocytes from 9-12-month-old *Wt* and *Sirt7^-/-^* samples. n = 3 mice per genotype and age group. **i)** Schematic diagram to illustrate the working model of how SIRT7 contributes to male mice fertility.

## DISCUSSION

In this study, we analyzed the effects of *Sirt7* deletion on mouse spermatogenesis. Our results demonstrate *Sirt7^-/-^* male mice exhibited an age-dependent subfertility characterized by early onset of defects typically occurring during natural aging in mice^23^. These defects included age-dependent decreased fecundity (fewer and smaller litters), increased histological abnormalities in seminiferous tubules, and impaired spermatogenesis. Further investigation revealed transcriptome changes that mimicked those of aged testes and constitutive H3K36ac epigenetic abnormalities in spermatogonia and spermatocytes.

Our work helps fill the gap of knowledge on the roles of sirtuins in spermatogenesis. Our analysis of publicly available transcriptomic data on human^33^ and mouse^34^ testes revealed distinct sirtuin expression patterns that suggest important roles during spermatogenesis. In human testes, *Sirt7* transcripts were the most abundant of all sirtuins in both spermatogonia and spermatocytes. In mouse, *Sirt7* was amongst the highest expressed sirtuins in undifferentiated spermatogonia together with *Sirt1*, *Sirt2* and *Sirt3* (Extended Data Fig. 4b-d), suggesting different requirements for sirtuin activity between humans and mice.

SIRT7 directly deacetylates H3K36ac and H3K18ac in somatic cells^26,29^. Although global levels of H3K18ac did not change upon SIRT7 deletion in testes, we observed a significant increase in H3K36ac regardless of age (Fig. 3). Consistent with H3K36ac being a *bona fide* substrate of SIRT7^29^, H3K36ac levels were inversely correlated with *Sirt7* transcript levels, being low in spermatogonia and early spermatocytes, including in leptonema and zygonema of prophase I, showing a mild increase during mid-pachynema and diplonema, and being high in round and elongating spermatids, where *Sirt7* transcript levels were the lowest (Fig. 4). Because previous work showed stage-specific changes in H3K27ac in male germ cells during aging^38^, our results suggest age-related histone acetylation alterations vary between different histone lysine residues.

In somatic cells, H3K36ac is prominently enriched at gene promoters and is proposed to modulate transcription^39^. Despite the dramatic increase in H3K36ac levels in *Sirt7^-/-^* testes, analyses of the transcriptomes did not indicate a preference for gene expression upregulation (Extended Data Fig. 2a), suggesting a role for this epigenetic mark outside of transcriptional regulation. Previous work in *Saccharomyces cerevisiae* established a positive correlation between H3K36ac and chromatin digestibility after genotoxic damage^30^. Our results also point to altered chromatin accessibility in the absence of SIRT7 in a spermatocyte-derived cell line (Fig. 5b, c), raising the possibility that *Sirt7^-/-^* spermatogonia and early spermatocytes also have compromised chromatin dynamics. Additionally, our data suggest H3K36ac influences nucleosome stability, but future experiments are necessary to define the molecular mechanisms by which this histone mark affects chromatin structure. Studies using human mesenchymal stem cells showed that SIRT7 promotes chromatin compaction through H3K9me3 regulation, indicating that SIRT7 may regulate chromatin accessibility through multiple epigenetic mechanisms^40^. Although our transcriptome analyses of 2-3-month-old *Sirt7^-/-^* testes revealed downregulation of the *Suv39h2* histone methyltransferase and upregulation of the *Kdm4d* histone demethylase (Extended Data Fig. 2b, c), we did not detect significant changes in global H3K9me3 levels in *Sirt7^-/-^* testes. Future investigation should resolve genome-wide maps of H3K36ac in germ cells and potential crosstalk with other epigenetic marks.

We found that testis aging in *Wt* animals is associated with increased DNA damage in undifferentiated spermatogonia, and this effect was exacerbated in *Sirt7^-/-^* testes (Fig. 6c, d). Since undifferentiated spermatogonia include the SSC pool^5^, which sustains steady-state spermatogenesis, the premature reproductive decline in *Sirt7^-/-^*males may be linked to defects in this population. Indeed, our findings align with previous studies in somatic tissues, which correlate stem cell genome damage accumulation with age-related physiological decline^41^. In support of this concept, SIRT7 is described to play key roles in hematopoietic^42^ and hair follicle^43^ stem cells, suggesting a broader role for SIRT7 in safeguarding stem cell homeostasis. SIRT7 deficiency also results in age-dependent spermatogenesis defects (Fig. 7) and DNA-damaged sperm (Fig. 1h- k), which may be directly linked to the observed genome instability in spermatogonia. However, further investigations are required to define additional SIRT7 roles at different developmental stages.

Reintroduction of SIRT7 into *Sirt7^-/-^* GC-2spd(ts) cells reduced γH2AX formation in the presence of external genotoxins (Fig. 5c, d and Extended Data Fig. 5), suggesting an important role of SIRT7 in genome damage prevention upon environmental stress. In somatic cells, SIRT7 participates in DSB repair through H3K18ac deacetylation^22^, which favors the recruitment of DNA repair factors at break sites. In yeast, H3K36ac favors homology recombination-mediated repair^30^ but whether it plays a similar role in mammalian cells is unknown. Our results indicated that SIRT7 deficiency increased H3K36ac (Figs. 3,4) and altered chromatin accessibility (Fig. 5), but further research is necessary to elucidate the impact of H3K36ac on DSB repair and genome maintenance.

Together, our results show SIRT7 is an important epigenetic factor in testicular aging regulation. Our findings indicate that SIRT7 has crucial roles in spermatogonia and spermatocytes, where it regulates H3K36ac and genome maintenance, thereby ensuring a sustained supply of high-quality gametes with age. Our results have important implications for potential interventions aimed at mitigating the age-related decline of male reproductive capacity.

## MATERIALS AND METHODS

### Animals and sample collection

The generation of *Sirt7^-/-^* mice on a 129/Sv genetic background has been previously described^22^. Animals were bred in the animal facilities of the Comparative Medicine and Bioimage Centre (CMCiB) of the Gemans Trias i Pujol Research Institute (IGTP) and Rutgers University, respectively. At both institutions, all animal procedures were approved by the Institutional Animal Care and Ethics. Experiments were performed on mice at 2-3, 4-6 and 9-12 months of age. Mice were humanely euthanized by CO_2_ inhalation and further cervical dislocation before the dissection of reproductive tissues. All animal experiments performed in this study were approved by the Rutgers IACUC (protocol #201702497) and the Catalan Government (protocol #10472).

### Fertility trials

For fertility assessments, *Wt* and *Sirt7^−/−^* single males were caged with either *Wt* outbred CF1 or *Wt* 129S1/Sv single females and co-housed until the end of the experiment. All animals were 8 weeks old when the study started. Total number of litters, pups per litter were recorded until seven litters were produced by control matings.

### Histological analyses

Testes were dissected, rinsed in 1X PBS, and fixed in Bouin’s Solution (Sigma-Aldrich, HT10132) overnight at 4°C. The following day, samples were washed and dehydrated before paraffin embedding using standard methods. Five µm sections of paraffin-embedded testis were stained with hematoxylin and periodic acid Schiff’s (PAS) stains using standard methods and coverslips mounted with DPX Mountant (Sigma-Aldrich, 06522). Images were acquired on an Olympus BX53 microscope equipped with an Olympus SC180 camera.

### Sperm comet assay

For the detection of DSBs, neutral comet assays were performed as previously described^16^, with technical adaptations for sperm. Briefly, sperm suspensions were spun for 4 min at 600 x g and pellets washed in 1X PBS and resuspended at a concentration of 2.5x10^6^ cells / ml. Ten µl of this suspension were diluted into 100 µl of 0.05% Low Melting Point (LMP) agarose (Lonza, #50081) and spread in a uniform layer on a Superfrost Plus glass slide (Epredia J1800AMNZ) previously coated with a layer of 1.75% agarose (Lonza, #50004). Cells were lysed in two steps. First, slides were incubated in lysis buffer (2.5 M NaCl, 100 mM EDTA, 10 mM Tris, 1% Sarkosyl, 1% Triton X-100, pH 10) supplemented with 40 mM DTT for 1 hr at room temperature (RT), and then in lysis buffer supplemented with 0.2 mg/ml Proteinase K (Apollo Scientific, BIP4205) at 37°C overnight. Next, slides were washed in TBE buffer (90 mM Tris, 90 mM Boric Acid, 2 mM EDTA, pH 8.5) and subsequently subjected to electrophoresis (0.6V / cm) in TBE for 25 min. Following electrophoresis, slides were stained with Hoechst (Thermo Scientific, 62249) before microscopic analyses. Comet images were captured on an Olympus BX51 fluorescence microscope equipped with an Olympus DP73 camera and analyzed using Comet Score software (TriTek). As recommended^44,45^, comets with ‘hedgehog morphology’, indicative of highly damaged DNA, were not included in the determination of % DNA in tail and were quantified separately.

### RNA isolation and sequencing

Testes were flash-frozen in liquid nitrogen immediately after euthanasia and stored at -80°C until all replicates were collected. Sample processing was performed at the same time for all replicates. The testis tunicae were removed, and a testis piece weighing <30 mg was sectioned for processing. The RNeasy Plus Mini kit (Qiagen #74134) was used for RNA isolation as per manufacturer’s instructions. Final RNA concentration was determined via NanoDrop (Thermo Fisher Scientific). RNA integrity was verified by running an RNA gel before proceeding to the next stage. Samples were sent to Novogene for processing through mRNA-seq pipeline. For analysis, datasets were first normalized by removing low count genes (<10 counts across all samples). Differentially expressed gene (DEG) analysis was performed with the DESeq2 package^46^. Pair-wise comparisons were calculated using the Wald test. Unsupervised hierarchical clustering was performed on significant DEGs (P adj < 0.1) after performing a likelihood ratio test across all samples. Gene Set Enrichment Analysis (GSEA) was performed using the Broad Institute GSEA software. DEG-obtained log_2_(FC) values were used as inputs for the GSEA. Molecular signatures were obtained from MSigDB (UC San Diego / Broad Institute) and Enrichr^47^.

### Publicly available RNA-seq dataset analyses

Sirtuin expression levels on the different testis cell populations were obtained from publicly available scRNA- seq data from young adult humans^48^ and from adult mice^34^. Processed data files from adult mice were obtained from Gene Omnibus Expression (GEO) accession code GSE112393. Expression plots from human datasets were retrieved from the public online resource ‘Human testis Atlas (https://humantestisatlas.shinyapps.io/humantestisatlas1/)^33^.

### cDNA synthesis and RT-qPCR assay

cDNA was synthesized from previously isolated RNA using a Transcriptor First Strand cDNA Synthesis Kit (Roche, 04379012001) following the manufacturer’s instructions. Real-time quantitative Polymerase Chain Reaction (RT-qPCR) was run with the QuantStudio 5 Real-Time PCR System (Thermo Fisher Scientific) using SYBR Green PCR Master Mix (Applied Biosystems, 4312704). Relative gene expression values were normalized to *actin beta (Actb)* levels. Oligonucleotide sequences are listed in Supplementary Table 1.

### CRISPR Cas9-mediated knock out of SIRT7 and clone generation

Guide crRNAs were designed using the IDT Alt-R CRISPR design tool (Supplementary Table 2). TracrRNA (IDT, 1072532), crRNAs and Cas9 nuclease (IDT, 1081058) were nucleofected into spermatocyte-derived GC- 2spd(ts) cells (ATCC CRL-2196) using a Nucleofector System (Lonza) following manufacturer’s protocols. Nucleofected single cells were sorted to generate clones. *Wt* clones were also generated by sorting single cells from the original pool. SIRT7 levels were assessed by western blotting and three clones of each genotype were chosen to perform experiments.

### MNase assay

Two million *Wt* and *Sirt7^-/-^* GC-2spd(ts) were pelleted, resuspended in 1 ml RSB buffer (10 mM Tris pH 7.8, 10 mM NaCl, 3 mM MgCl2, 1% NP-40, 0.5 mM DTT, 1X Complete Protease Inhibitor (Roche, 11836170001)) and incubated for 10 min at 4 °C. Nuclei were pelleted by centrifuging 30 sec at maximum speed and resuspended in 400 μl of nuclear buffer (20 mM KCl, 20 mM Tris pH 8, 70 mM NaCl, 3 mM CaCl2, 1X Complete Protease Inhibitor). Thirty units of micrococcal nuclease (MNase) (Thermo Scientific, 88216) were added to the suspension and incubated at RT. To stop the reaction at each digestion time point, 50 μl of the suspension were retrieved, mixed with 3 μl EDTA 0.5 μM and cooled on ice. Digested DNA was purified, quantified and loaded on a 1.8% agarose gel for electrophoresis and analysis.

### Cell transfections and treatments

*Wt* and *Sirt7^-/-^* GC-2spd(ts) (ATCC CRL-2196) were cultured in Dulbecco’s modified Eagle’s medium (DMEM) (Gibco, 31966-021) supplemented with 10% FBS (Gibco, 10270-106) and 1% PenStrep (BioWest, l0022-100) at 37°C, 5% CO_2_. Cells were transiently transfected with empty vector (EV) or SIRT7 pcDNA4T0 plasmids^22^ using 4 μg polyethylenimine (Polysciences, 23966) per μg of DNA, harvested 48 hr post-transfection.

For genotoxic treatments, culture media was supplemented with 0.1 µM doxorubicin (DOX) (Shelleckchem, S1208). In the case of transfected cells, treatments were added 24 hr after transfection. In all cases, 24 hr after treatment cells were washed and pelleted for protein extraction.

### Protein extraction

To obtain protein lysates, testes were dissected, snap-frozen in liquid nitrogen, and stored at -80°C until processing. Frozen testes were mechanically homogenized using a Polytron homogenizer (Inycom, PT 1200 E) in cold RIPA buffer (50 mM Tris pH 7.8, 150 mM NaCl, 0.5% Deoxycholic acid, 0.1% SDS, 1% NP-40, 1X Complete Protease Inhibitor (Roche, 11836170001)) and sonicated. For GC-2spd(ts) protein lysates, cellular pellets were lysed in cold RIPA buffer and sonicated.

### SDS-PAGE and western blot

Protein lysates were mixed with 5X Laemmli sample buffer (2% SDS, 10% glycerol, 60 mM Tris pH 6.8, 0.01% bromophenol blue) supplemented with 10% 2–mercaptoethanol (Sigma-Aldrich, M3148), and boiled at 95°C for 5 min. Protein extracts were separated by SDS-PAGE (15% polyacrylamide) and then transferred to nitrocellulose membrane in transfer buffer (200 mM glycine, 25 mM Tris-HCl, 0.1% SDS, 20% methanol). Membranes were blocked with 5% (w/v) non-fat milk (in 1X PBS, 0.1% Triton X-100), and then incubated with primary antibodies (Supplementary Table 3) diluted in the same blocking solution overnight at 4℃. Membranes were washed three times for 10 min with 1X PBS + 0.1% Tween-20 and incubated 1 hr at RT with secondary antibodies (Supplementary Table 3). After washing the membranes thrice for 10 min, antibody binding was visualized with an ECL chemiluminescence system (Thermo Fisher Scientific, SuperSignal detection kit) and exposure of the membrane to iBright Imaging Systems (Thermo Fisher Scientific).

### Immunostaining of testicular sections

Testes were dissected and fixed overnight in 4% paraformaldehyde (PFA) at 4 °C. After three 15-min washes in H_2_O, samples were incubated overnight in 30% sucrose. Then, testes were embedded in O.C.T. Compound (Tissue Tek, 4583) and frozen in dry ice. Five µm sections were cut with a cryostat, placed in a Superfrost Plus slide (Epredia, J1800AMNZ) and kept at –80 °C until use. Antigen retrieval was performed in 10 mM sodium citrate, 0.05% Tween-20, pH 6 at 95°C during 20 min. Slides were allowed to cool before permeabilization in 0.1% TritonX-100, 1X PBS for 10 min and further blocking in blocking buffer (0.3% BSA, 0.03% glycine, 0.01% Tween-20, 0.02% TritonX-100, 1X PBS) for 1 hr. Primary antibodies (Supplementary Table 4) were diluted in blocking buffer and incubated overnight at 4°C. Secondary antibodies (Supplementary Table 4) were incubated 1 hr at 37 °C. Samples were counterstained with DAPI and mounted with Vectashield VIBRANCE mounting media (VectorLabs). Images were acquired with a STELLARIS 8 confocal microscope (Leica). Image analyses were performed using FIJI^49^.

### Preparation of meiotic chromosome spreads

Meiotic chromosome spreads were prepared as previously described^50^, with minor modifications. Briefly, 5x5 mm pieces of frozen testes were minced on a cold surface with a scalpel to obtain a homogeneous cell suspension. Cells were resuspended in 200 µl cold 1X PBS, and 1X Complete Protease Inhibitor (Roche, 11836170001). Twenty-five µl of cell suspension were placed on Superfrost Plus slides and 80 µl of 1% Lipsol detergent solution were added and distributed along the slide surface. Slides were incubated in a humid chamber for 17 min at RT. Next, 150 µl of fixative solution (1% paraformaldehyde (PFA), 0.15% Triton X-100, 1X Complete Protease Inhibitor (Roche, 11836170001), pH 9.3) were added to the slides. Samples were fixed in the humid chamber for 2 hr at RT and air-dried for 30 min. Finally, slides were washed twice for 2 min in 0.4% Photo-Flo (Kodak) in 1X PBS and twice for 2 min in 0.4% Photo-Flo in H_2_O. Slides were air-dried and stored at -80°C until use.

### Immunostaining of meitoic chromosome spreads

Immunostaining of chromosome spreads was performed as previously described^21^. Briefly, slides were blocked for 40 min in blocking buffer (1X PBS, 0.5% BSA 0.1%, Tween-20). Primary antibodies (Supplementary Table 4) were diluted in blocking buffer and incubated in a humid chamber overnight at 4°C. Slides were washed thrice for 15 min in 1X PBS, 0.1% Tween and incubated with secondary antibodies (Supplementary Table 4) diluted in blocking buffer for 1 hr at 37°C. After three 15 min washes, slides were counterstained with DAPI and mounted with Vectashield VIBRANCE mounting media (VectorLabs). Images were acquired with a STELLARIS 8 confocal microscope (Leica). Image analyses were performed using FIJI^49^.

### Statistical analyses

Results are presented as the mean ± standard error (SEM) or density plots, with P-values calculated by Student’s t-test, one-factor ANOVA or two-factor ANOVA (significance set at P-values < *0.05, **0.01, ***0.001,

****0.0001), according to the particularities of each dataset.

## Acknowledgements

This work was supported by FI predoctoral fellowship by the Catalan Government (to AGS; #2022 FI_B 00924), UAB PPC-2024 grant (to BNV), a Pilot Project Funding from the American Society of Reproductive Medicine (to KS), NIH grants R21HD105963 (to KS and CG), R01HD110170 (to CG), MICIU/AEI/10.13039/501100011033 and FEDER-EU grant PID2023-148760OB-I00 (to AV) and PID2022-138905OB-I00 (to IR), and FEDER-EU 2017-SGR-148 and 2021-SGR-01378 grants from the Catalan Government Agency AGAUR (to AV).

## DATA AVAILABILITY

The bulk RNA-seq raw and processed data generated in this study have been deposited in the GEO under the accession number GSE291149.

## EXTENDED DATA

**Extended Data Figure 1:**
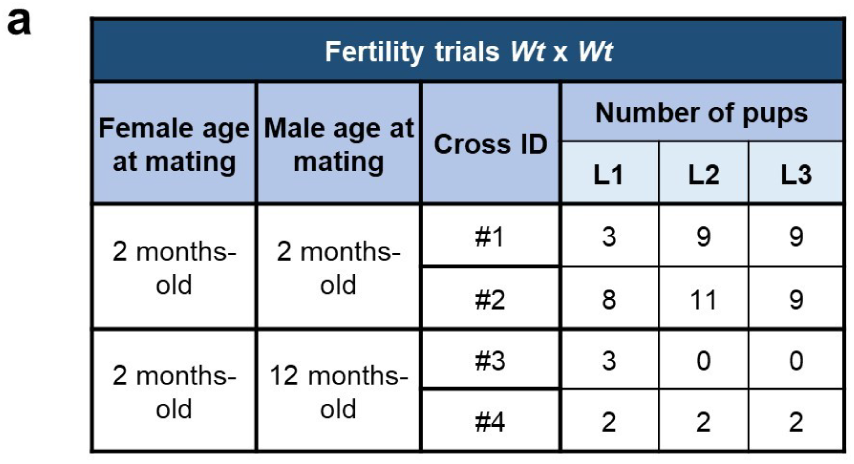
Aging fertility trials. **a)** Pups per litter sired by *Wt* x *Wt* matings with males of different ages. n = 2 matings per age group. 3 litters were recorded for each mating when possible.

**Extended Data Figure 2:**
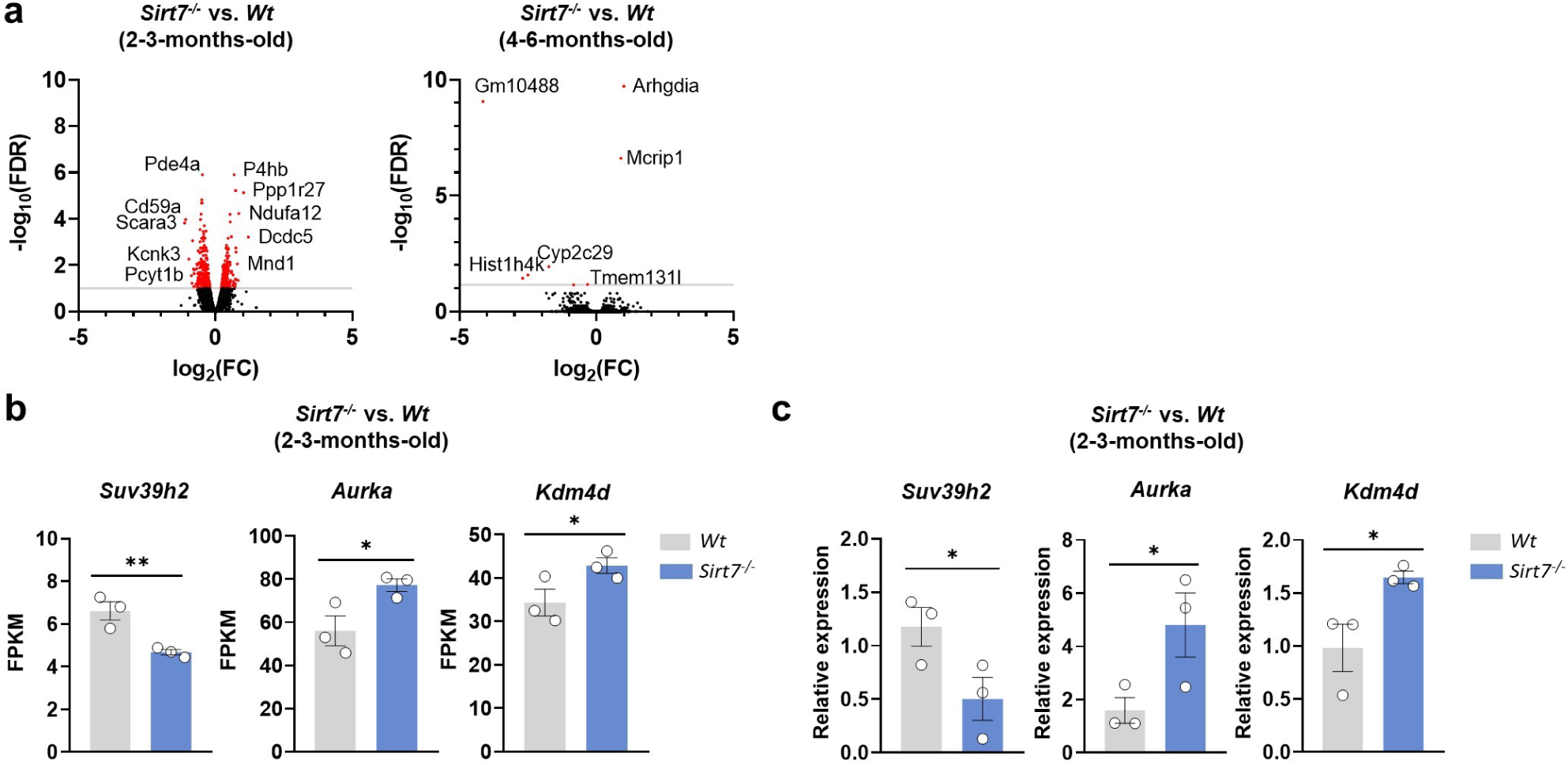
Early transcriptomic deregulation due to lack of SIRT7. **a)** Volcano plots derived from RNA-seq data from 2-3- and 4-6-month-old testis showing the DEGs between *Sirt7^-/-^* and *Wt* samples. Significant DEGs (P adj < 0.1) are shown in red. **b)** FPKM values of the differentially expressed chromatin modifiers *Suv39h2*, *Aurka* and *Kdm4d* in 2-3-month-old *Sirt7^-/-^* samples. **c)** Validation of DEGs found in RNA-seq data by RT-qPCR in 2-3-month-old *Wt* and *Sirt7^-/-^* testis. n = 3 samples per group.

**Extended Data Figure 3:**
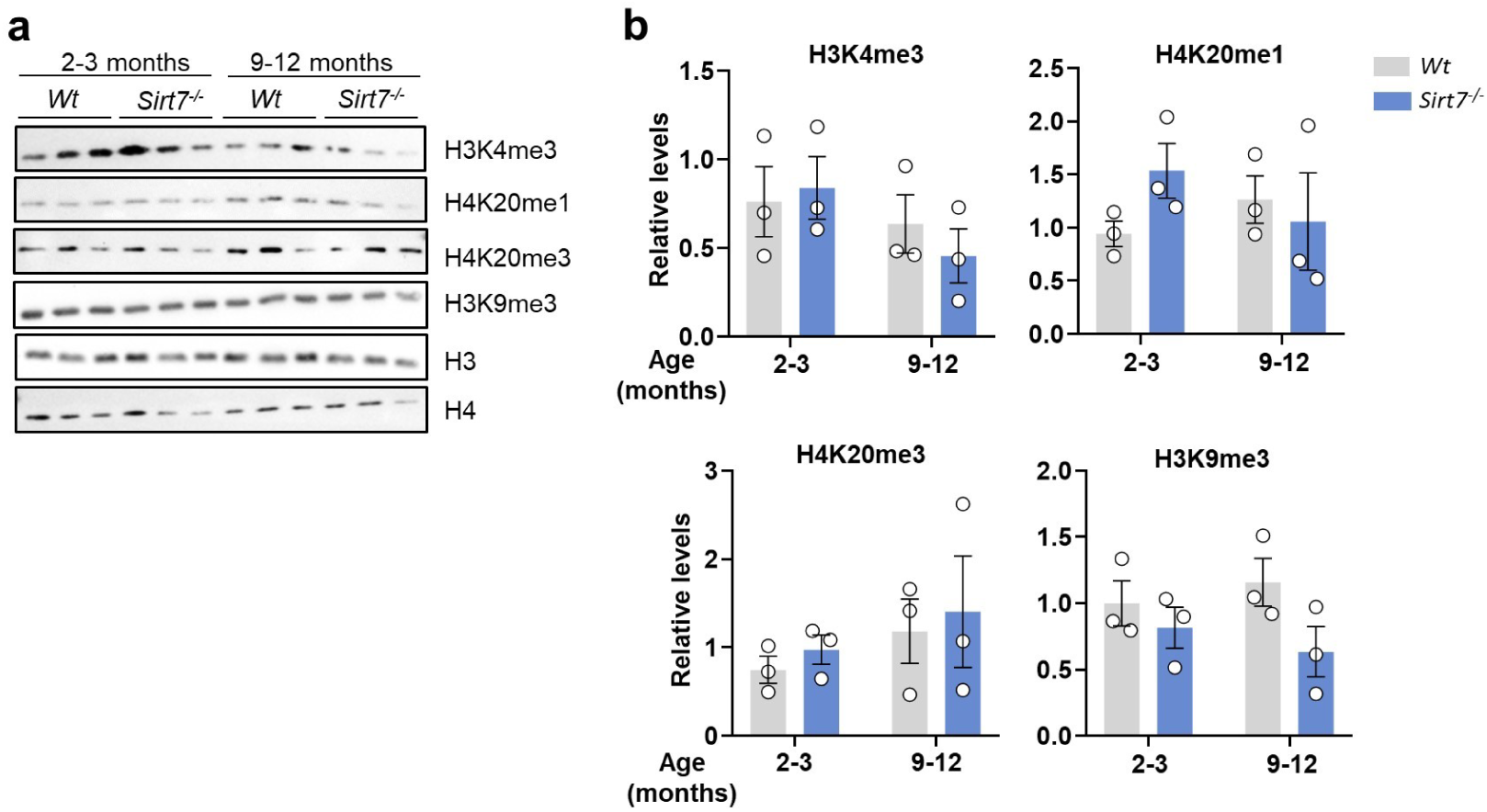
Spermatogenesis and Sirtuin linked epigenetic marks in *Wt* an *Sirt7^-/-^* samples. **a, b)** Western blot (a) and quantifications (b) of epigenetic marks in 2-3- and 9-12-month-old *Wt* and *Sirt7^-/-^* testes. Densitometry-based quantification of the levels of each epigenetic mark were normalized to H3 or H4 levels. n = 3 testis samples per genotype and per age group.

**Extended Data Figure 4:**
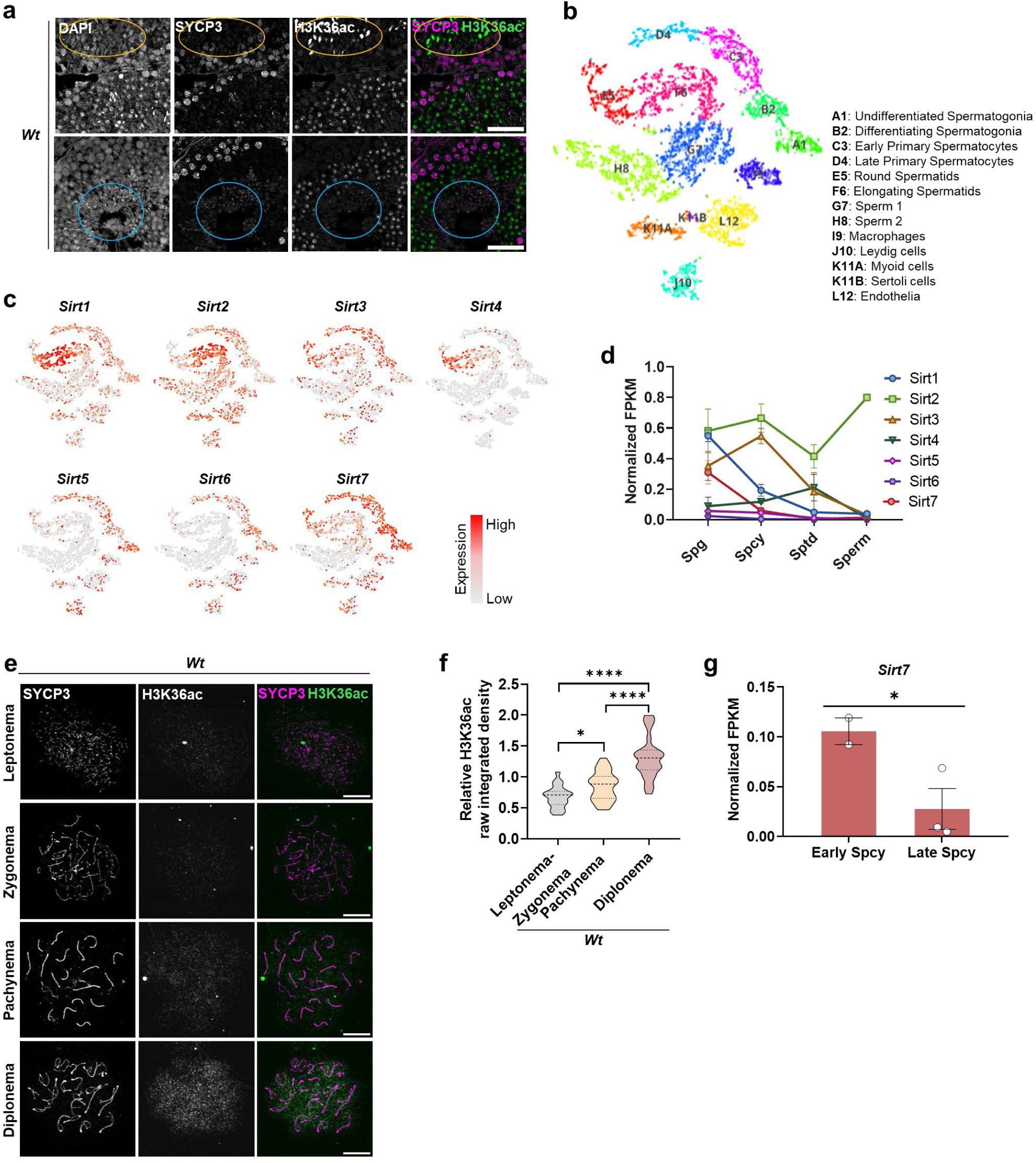
SIRT7 levels in germ cells correlate to H3K36ac levels and change with aging. **a)** Representative immunostainings of SYCP3 and H3K36ac in 2-3-month-old *Wt* testis sections. Orange circles marks elongating spermatids and blue circles elongated spermatids. **b)** tSNE plot of single-cell transcriptome data from young human testes showing all testis cell groups. **c)** tSNE plots with sirtuin expression pattern in the different testis populations shown in (b). **d)** Data from scRNA-seq of mouse testes showing the expression levels of mouse sirtuins during spermatogenesis. Spg, Spermatogonia; Spermatocytes, Spcy; Spermatids, Sptd. **e)** Representative immunostaining images of SYCP3 and H3K36ac-stained prophase I cells in spermatocyte chromosome spreads of *Wt* samples. **f)** Quantification of H3K36ac immunofluorescence intensity in meiotic prophase I stages. **g)** Data from scRNA-seq of mouse testes showing the differential expression of *Sirt7* between early and late spermatocytes.

**Extended Data Figure 5:**
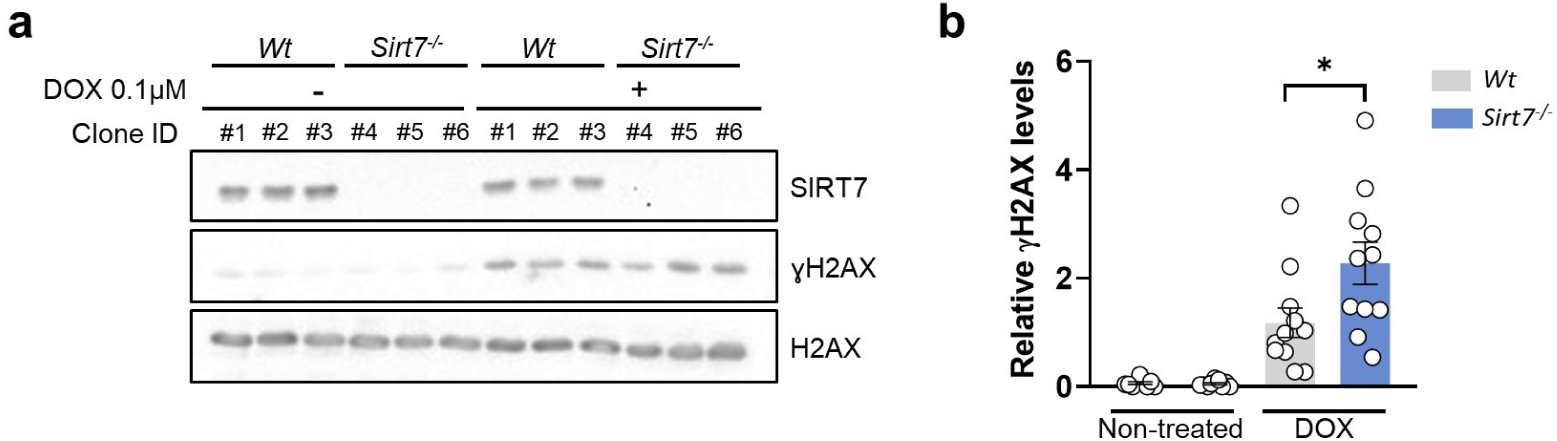
DNA damage after genotoxic stress in *Wt* and *Sirt7^-/-^* GC-2spd(ts). **a)** Western blot of SIRT7 and ɣH2AX in *Wt* and *Sirt7^-/-^*GC-2spd(ts) in non-treated and 0.1 µM DOX-treated *Sirt7^-/-^* GC-2spd(ts) in 3 independent clones per gentoype. n = 4 experiments. **b)** Densitometry-based quantification of ɣH2AX levels of (a) relative to H2AX levels.

**Extended Data Figure 6.**
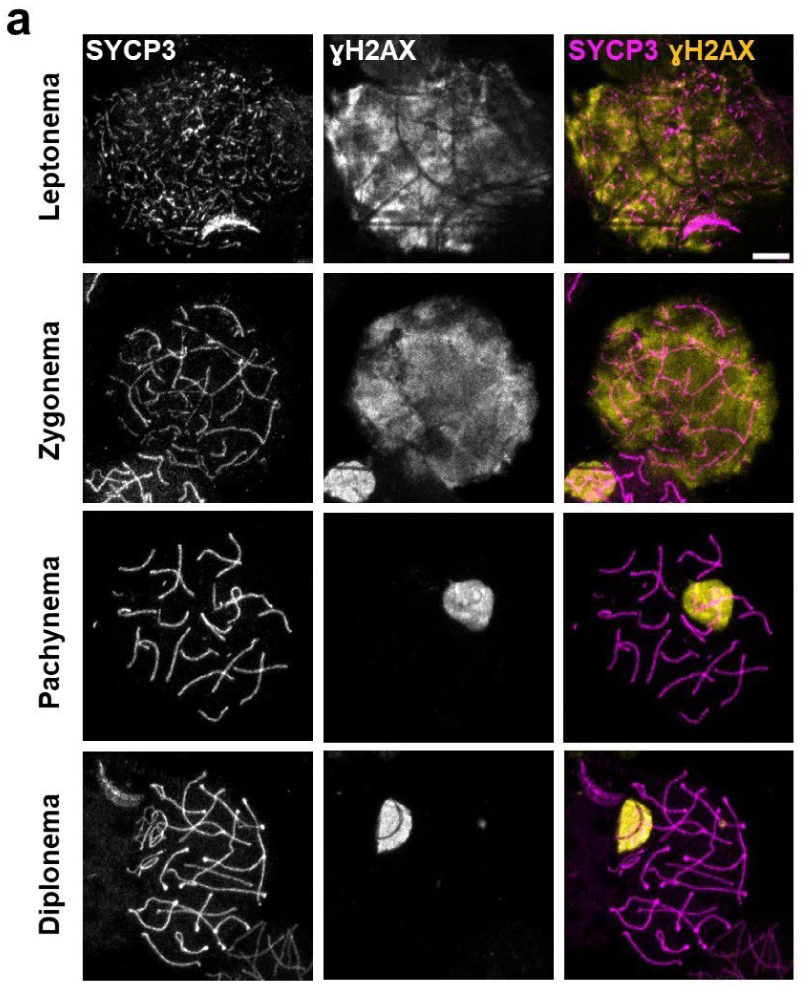
ƔH2AX in meiotic prophase I spermatocytes. **a)** Representative immunostaining images of the meiotic prophase I stages after staining for SYCP3 and ɣH2AX. Scale bar = 10 µm.

## SUPPLEMENTARY INFORMATION

**Supplementary Table 1:**
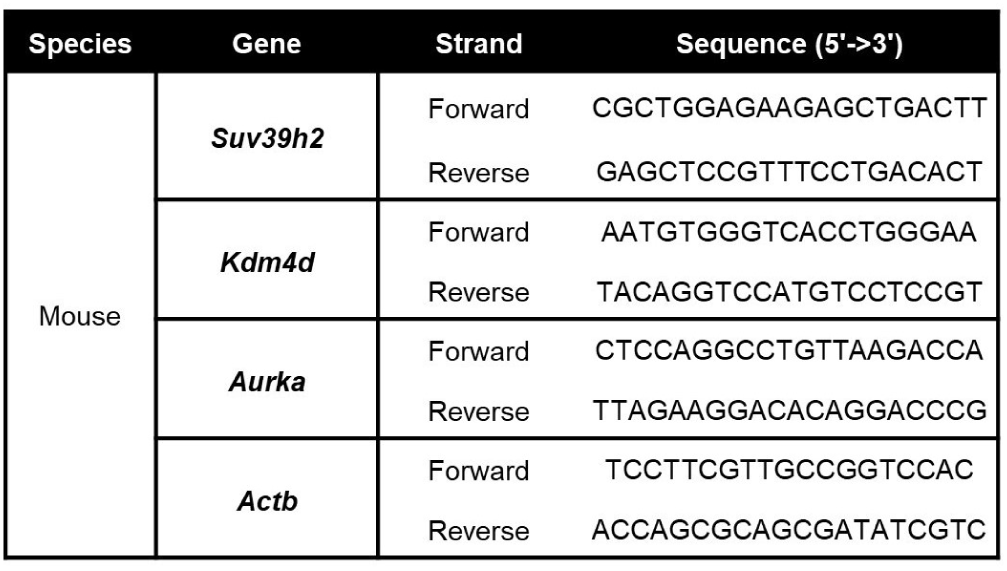
RT-qPCR primer list.

**Supplementary Table 2:** Sirt7 knock-out CRISPR crRNA guide sequences.

| Gene | Guide | Exon | Sequence (5'→3') |
| --- | --- | --- | --- |
| *Sirt7* | 1 | 1 | /AltR1/rCrCrA rUrUrA rGrGrA rCrCrC rCrGrA rUrArA rUrCrG rUrUrU rUrArG rArGrC rUrArU rGrCrU /AltR2/ |
|  | 2 | 1 | /AltR1/rCrGrG rArGrC rGrCrA rArArG rCrUrG rCrUrG rArGrG rUrUrU rUrArG rArGrC rUrArU rGrCrU /AltR2/ |

**Supplementary Table 3:** List of antibodies for Western Blot.

| Primary antibodies | Host | Dilution | Source | Catalogue number |
| --- | --- | --- | --- | --- |
| H3K36ac (D9T5Q) | Rabbit | 1:1000 | Cell Signaling Technology | 27683 |
| H3K9me3 | Rabbit | 1:1000 | Abcam LTD. | ab8898 |
| H3K18ac | Rabbit | 1:1000 | Abcam LTD. | ab1191 |
| H3K36me2 | Rabbit | 1:1000 | Cell Signaling Technology | C75H12 |
| H3K36me3 (D5A7) | Rabbit | 1:1000 | Cell Signaling Technology | 4909S |
| H3K4me3 | Rabbit | 1:1000 | Abcam LTD. | ab8580 |
| H4K20me1 | Rabbit | 1:1000 | Abcam LTD. | ab9051 |
| H4K20me3 | Rabbit | 1:1000 | Novus Biologicals | 5E10-D8 |
| γH2AX | Rabbit | 1:500 | Abcam LTD. | ab2893 |
| H3 | Rabbit | 1:10.000 | Abcam LTD. | ab1791 |
| H4 | Rabbit | 1:1000 | Cell Signaling Technology | 2935 |
| H2AX | Rabbit | 1:1000 | Abcam LTD. | ab11175 |
| SIRT7 | Mouse | 1:1000 | Santa Cruz Biotechnology | sc-365344 |
| Secondary antibodies | Host | Dilution | Source | Catalogue number |
| Anti-Mouse IgG-HRP | Rabbit | 1:10.000 | Sigma-Aldrich | A9044 |
| Anti-Rabbit IgG- HRP | Goat | 1:10.000 | Sigma-Aldrich | A0545 |

**Supplementary Table 4.** List of antibodies for immunostaining.

| Primary antibodies | Host species | Dilution for IF in chromosome spreads | Dilution for IF in testicular sections | Source | Catalogue number |
| --- | --- | --- | --- | --- | --- |
| SYCP3 | Mouse | 1:500 | 1:200 | Santa Cruz Biotechnology | sc-74569 |
| SYCP3 | Rabbit | 1:500 | - | Novus Biologicals | NB300-232 |
| γH2AX | Mouse | 1:2000 | 1:1000 | Sigma Aldrich | 05-636-I |
| RAD51 | Rabbit | 1:250 | - | Sigma Aldrich | PC-130 |
| HORMAD1 | Rabbit | 1:200 | - | Proteintech | 13917-1-AP |
| H3K36ac | Rabbit | 1:1000 | 1:500 | Cell Signaling Technology | 27683 |
| PLZF | Rabbit | - | 1:1000 | Abcam LTD. | ab189849 |
| Secondary antibodies | Host species | Dilution for IF in chromosome spreads | Dilution for IF in testicular sections | Source | Catalogue number |
| Alexa Fluor-488 anti-rabbit IgG | Goat | 1:500 | 1:500 | Thermo Fisher | A-11034 |
| Alexa Fluor-555 anti-mouse IgG | Goat | 1:500 | 1:500 | Thermo Fisher | A-32727 |
| Alexa Fluor-647 anti-rabbit IgG | Goat | 1:500 | 1:500 | Thermo Fisher | A-21245 |

## Notes

### Competing Interest Statement

The authors have declared no competing interest.

